# A population genomics approach to assessing the genetic basis of within-host microevolution underlying recurrent cryptococcal meningitis infection

**DOI:** 10.1101/083469

**Authors:** Johanna Rhodes, Mathew A. Beale, Mathieu Vanhove, Joseph N. Jarvis, Shichina Kannambath, John A. Simpson, Anthea Ryan, Graeme Meintjes, Thomas S. Harrison, Matthew C. Fisher, Tihana Bicanic

## Abstract

Recurrence of meningitis due to *Cryptococcus neoformans* after treatment causes substantial mortality in HIV/AIDS patients across sub-Saharan Africa. In order to determine whether recurrence occurred due to relapse of the original infecting isolate or reinfection with a different isolate weeks or months after initial treatment, we used whole-genome sequencing to assess the genetic basis of infection in 17 HIV-infected individuals with recurrent cryptococcal meningitis. Comparisons revealed a clonal relationship for 15 pairs of isolates recovered before and after recurrence showing relapse of the original infection. The two remaining pairs showed high levels of genetic heterogeneity; in one pair we found this to be a result of infection by mixed genotypes, whilst the second was a result of nonsense mutations in the gene encoding the DNA mismatch repair proteins *MSH2, MSH5* and *RAD5*. These nonsense mutations led to a hypermutator state, leading to dramatically elevated rates of synonymous and non-synonymous substitutions. Hypermutator phenotypes owing to nonsense mutations in these genes have not previously been reported in *Cryptococcus neoformans* and represent a novel pathway for rapid within-host adaptation and evolution of resistance to firstline antifungal drugs.

## Introduction

The HIV/AIDS pandemic has led to a large population of profoundly immunocompromised individuals that are vulnerable to infection by the opportunistic fungus pathogen *Cryptococcus neoformans* (*Cn*) (1). This mycosis poses a considerable public health problem in sub-Saharan Africa, which has the highest estimated annual incidence of cryptococcal meningitis (CM) globally (2), with the majority of infections caused by *Cryptococcus neoformans sensu stricto* (previously referred to as *Cryptococcus neoformans* var. *grubii*) (3).

Standard treatment for HIV-associated CM includes the long-term use of azole drugs such as fluconazole, following initial 1-2 week induction treatment with amphotericin B, which is often not available (4). Microevolution occurs in response to drug pressure, leading to resistance, a phenomenon previously described in *Cn* (5, 6). Patients who appear successfully treated (evidenced by symptom resolution and sterilisation of cerebral spinal fluid (CSF)) can relapse due to persisting infections, which in some cases appear to have evolved resistance to firstline antifungal drugs. In the absence of continued antifungal therapy and restoration of their immune system through antiretroviral therapy (ART), patients with HIV/AIDS also have a high probability of recurrence of CM (7).

Various methods of within-host evolution are available to eukaryotic pathogens, most notably sexual and parasexual reproduction, although these are difficult to observe in fungi due to being cryptic. Aneuploidy, recombination in the telomeres, and mutator states (8) also provide means of rapid within-host evolution, with other mechanisms still likely to be discovered. The accumulation of SNPs alongside copy number variation and aneuploidy has been witnessed during infection in different fungal pathogen species, enabling rapid adaptive evolution (9) and conferring resistance to antifungal dugs (10). Candidiasis is caused by numerous *Candida* species, yet within-host evolution to these species differs. Infection by *Candida albicans* is usually susceptible to azole antifungal drugs, but resistance can evolve via the acquisition of drug-resistant aneuploid isolates, which contain an isochromosome of the left arm of chromosome 5 (11). The left arm of chromosome 5 contains two important genes involved in resistance to antifungals: *ERG11*, a target of azoles, and *TAC1*, a transcription factor that activates drug efflux pump expression. Conversely, *Candida glabrata* are intrinsically poorly susceptible to azoles, and have more recently evolved multi-drug resistance to both azoles and echinocandins (12-14).

Within-host diversity and recombination has been witnessed in eukaryotic pathogens, notably *C. albicans*: mutation and recombination rates can be increased under stressful conditions, such as drug treatment (15, 16), resulting in loss-of-heterozygosity (LOH) and aneuploidy (15). These genetic alterations contribute to the maintenance of a population of *C. albicans* within the host environment (15), and drug pressure can result in diverging levels of fitness (17).

Similar responses to antifungal drugs have been observed in *Cn*; Point mutations in the *Cn* ortholog *ERG11* were also shown to confer fluconazole resistance, by causing the amino acid substitution G484S (8). Sionov *et al*. (6) demonstrated large scale chromosomal duplications (primarily chromosome 1) are fundamental to overcoming fluconazole (FLC) drug pressure in a mouse model, contributing to failure of FLC therapy. The duplication of chromosome 1 included increased copy number of genes *ERG11*, the target of FLC, and *AFR1*, a transporter of azoles (6), although other genes are also thought to be involved in FLC resistance (18, 19). Previous studies of serially collected *Cn* isolates have confirmed in-host microevolution, including the occurrence of large-scale genomic rearrangements (20-22). Like *C. albicans*, the *Cn* genome is capable of undergoing chromosomal duplication and loss under stresses such as drug pressure or invasion of the human host (23). These chromosomal duplications are often lost when the selective pressure is removed (6).

The development of mutator states via hypermutability is a rapidly expanding area of study in bacteria, particularly *Pseudomonas aeruginosa* in cystic fibrosis (CF) patients. Here, hypermutability has been shown to have an association with antimicrobial resistance (24, 25), causing significant implications in the early treatment of cystic fibrosis patients to prevent chronic infection (25, 26). Few studies have explored hypermutation in pathogenic fungi; however, mutations in the yeast *Saccharomyces cerevisiae* genes *PMS1, MLH1* and *MSH2*, which are all involved in mismatch repair, have been shown to lead to 100- to 700- fold increases in mutations throughout the genome (27). Frameshift mutations in an ortholog of the mismatch repair gene *MSH* have also been shown to contribute to microevolution in the sister species of *Cn, Cryptococcus gattii* (28).

Here we describe a comparative genome-sequencing based approach to investigate microevolution in serially collected isolates of *Cn*. These isolates were grown and stored from fresh CSF of patients with CM, prior to starting and during antifungal therapy using induction with amphotericin B-based regimens, followed by fluconazole. We used whole-genome sequencing to describe the nature of infection in 17 patients to gain insights into the dynamics of recurrent infections.

## Materials and Methods

### Samples and patients

Sixteen South African patients and one Ugandan patient demonstrating clinical evidence of cryptococcal meningitis were studied. All patients were either part of observational studies or clinical trial (29-33). Ethical approval was obtained from the Wandsworth Research Ethics Committee covering St. George's University of London (29-32). In South Africa additional ethical approval was obtained from the University of Cape Town Research Ethics Committee; in Uganda, from the Research Ethics Committee of Mbarara University of Science and Technology. All patients initially presented with cryptococcal meningitis and were treated using induction therapy with 7-14 days' amphotericin B deoxycholate 0.7-1 mg/kg/d, with or without 100 mg/kg/d of flucytosine (with one patient, IFNR63, also receiving adjunctive interferon gamma), followed by fluconazole consolidation at 400 mg/d for 8 weeks and maintenance therapy at 200 mg/d for 6-months (*n* = 16 pairs), until immune restoration on ART with a CD4 count of > μ L. The single patient in Uganda received induction therapy with fluconazole mg/d for 2 weeks followed by fluconazole consolidation and maintenance and ART, as above (*n* = 1 pair). As part of study procedure, patients enrolled in clinical trials had quantitative cryptococcal cultures performed on serial CSF samples. Patients with a recurrence of their cryptococcal disease following initial treatment and positive CSF culture for *Cryptococcus* at the time of disease recurrence were included in the study. We studied the clinical cryptococcal isolates taken on initial diagnosis (prior to initiation of treatment) and compared each with the *Cryptococcus* isolated from CSF on recurrence of disease in the same patient (Table 1).

**Table 1:** Details of *Cn* isolates and MICs (if available) at time of isolation from South Africa and Uganda (Pair 7 only) used in this study. AmB = amphotericin B 1 mg/kg/d, as per hospital guidelines at that time, unless otherwise stated; VOR = voriconazole mg/d; 5FC = flucytosine; FLC = fluconazole.

| Pair # | Isolate ID | Day of isolation | Fluconazole MIC at isolation | Treatment of CM episode (if known) |
| --- | --- | --- | --- | --- |
| 1 | CCTP27 | 1 | 4 | AmB 3 days |
| 1 | CCTP27-d121 | 121 | 64 | VOR until CD4 > 200 cells/uL |
| 2 | CCTP32 | 1 | 4 | AmB 7 days |
| 2 | CCTP32-d132 | 132 | 6 | AmB 7 days |
| 3 | CCTP50 | 1 | 16 | AmB 14 days |
| 3 | CCTP50-d257 | 257 | 256 | AmB 14 days |
| 3 | CCTP50- | 409 | n/a | AmB 7 days |
|  | d409 |  |  |  |
| 4 | CCTP52 | 1 | n/a | FLC 400 mg/d |
| 4 | CCTP52-d55 | 55 | n/a | FLC 400 mg/d |
| 5 | RCT9 | 1 | 2 | AmB 0.7 mg/kg/d plus<br>5FC for 14 days |
| 5 | RCT9-d99 | 99 | n/a | AmB until death |
| 6 | RCT24 | 1 | 4 | AmB plus 5FC for 14<br>days |
| 6 | RCT24-d154 | 154 | 12 | FLC 800 mg/d |
| 7 | 1600-1 | 1 | n/a | FLC 16000 mg/d for 14<br>days |
| 7 | 1600-1-d106 | 106 | n/a |  |
| 8 | IFNR63 | 1 | n/a | AmB plus 5FC |
| 8 | IFNR63-d128 | 128 | n/a |  |
| 9 | IFNR24 | 1 | n/a | AmB 7 days |
| 9 | IFNR24-d101 | 101 | n/a |  |
| 10 | IFNR18 | 1 | n/a | AmB 7 days |
| 10 | IFNR18-d134 | 134 | n/a |  |
| 11 | IFNR14 | 1 | n/a | AmB 7 days |
| 11 | IFNR14-d97 | 97 | n/a |  |
| 12 | IFNR13 | 1 | n/a | AmB 7 days |
| 12 | IFNR13-d95 | 95 | 1 |  |
| 13 | IFNR6 | 1 | n/a | AmB 7 days |
| 13 | IFNR6-d73 | 73 | 1 |  |
| 14 | IFNR19 | 1 | n/a | AmB 7 days |
| 14 | IFNR19-d111 | 111 | n/a |  |
| 15 | IFNR11 | 1 | n/a | AmB 7 days |
| 15 | IFNR11-d203 | 203 | 12 |  |
| 16 | IFNR27 | 1 | n/a | AmB 7 days |
| 16 | IFNR27-d204 | 204 | n/a |  |
| 17 | IFNR23 | 1 | n/a | AmB 7 days |
| 17 | IFNR23-d179 | 179 | n/a |  |

**Table 2:** A high number of shared SNPs in most pairs indicate a shared common ancestor. Number of SNPs common to both initial and recurrent infection, along with number of SNPs and non-synonymous SNPs unique to each timepoint. Percentages given to 2 d.p.

| Pair | Common SNPs<br>(% total) | Day 1 SNPs<br>(% total) | No. Day<br>1<br>nsSNPs<br>(genes<br>mapped<br>) | Relapse SNPs<br>(% total) | No. Relapse<br>nsSNPs<br>(genes<br>mapped) | Relapse #2<br>SNPs (% total) | No. Relapse<br>#2 nsSNPs<br>(genes<br>mapped) |
| --- | --- | --- | --- | --- | --- | --- | --- |
| 1 | 13490 (98.49) | 97 (0.71) | 9 (8) | 110 (0.81) | 19 (8) |  |  |
| 2 | 289557<br>(99.29) | 1080 (0.37) | 110 (55) | 991 (0.34) | 108 (45) |  |  |
| 3 | 261718<br>(46.77) | 127631<br>(32.78) | 25331<br>(2741) | 45769 (14.88) | 8342 (2099) | 124498<br>(32.24) | 23957<br>(2541) |
| 4 | 289574<br>(99.22) | 1169 (0.40) | 145 (58) | 1096 (0.38) | 145 (56) |  |  |
| 5 | 289367<br>(99.20) | 1261 (0.43) | 127 (54) | 1080 (0.37) | 127 (59) |  |  |
| 6 | 13122 (95.78) | 133 (1.00) | 5 (2) | 445 (3.28) | 82 (56) |  |  |
| 7 | 47833 (97.83) | 522 (1.08) | 81 (65) | 537 (1.11) | 94 (72) |  |  |
| 8 | 46883 (98.76) | 294 (0.62) | 29 (9) | 296 (0.63) | 26 (16) |  |  |
| 9 | 289458<br>(99.27) | 1033 (0.36) | 141 (58) | 1109 (0.38) | 105 (55) |  |  |
| 10 | 12687 (98.29) | 104 (0.81) | 4 (3) | 117 (0.91) | 11 (5) |  |  |
| 11 | 12732 (98.38) | 110 (0.86) | 10 (2) | 99 (0.77) | 6 (5) |  |  |
| 12 | 221973<br>(99.37) | 636 (0.29) | 55 (27) | 778 (0.35) | 61 (30) |  |  |
| 13 | 48062 (98.58) | 396 (0.82) | 31 (11) | 294 (0.61) | 28 (15) |  |  |
| 14 | 28960 (95.66) | 288 (0.98) | 44 (19) | 1026 (3.42) | 105 (57) |  |  |
| 15 | 29736 (98.40) | 254 (0.85) | 21 (14) | 228 (0.76) | 17 (13) |  |  |
| 16 | 45437 (98.56) | 325 (0.71) | 30 (18) | 341 (0.74) | 30 (14) |  |  |
| 17 | 376568<br>(56.43) | 117436<br>(23.77) | 22567<br>(4153) | 173359<br>(31.52) | 33543<br>(4427) |  |  |

### Multi-locus sequence typing

To discern whether mixed or single genotype infections were extracted from CSF, multi-locus sequence typing (MLST) was performed on three independent colonies for a subset of the study isolates, according to the methods outlines in Meyer *et al*. (34), with modifications as outlined in Beale *et al*. (35).

### Molecular methods

*Cn* was isolated from HIV infected individuals on location by plating CSF onto Sabourand Dextrose (SD) agar (Oxoid, Fisher Scientific), and growing at 30°C for 48 hours. A representative sample of the *Cn* population was taken by selecting a broad ‘sweep’ of all colonies on the SD agar plate, which was stored in cryopreservative medium (80% SD broth, 20% glycerol) at -80°C until further testing. This approach ensures all genetic diversity is maintained through the process, and single colony picking only occurs at the final stage of liquid culture and DNA extraction.

Cn isolates from patient CSF were stored in 20% glycerol at -80°C until ready for testing. Frozen stocks were plated onto SD agar and cultured for 72 hours. A single colony was inoculated into 6ml Yeast Peptone Digest broth (Oxoid) supplemented with 0.5M NaCl and cultured at 37°C with agitation (165 rpm) for 40 hours, followed by genomic DNA extraction using the Masterpure Yeast DNA purification kit (Epicentre) modified by addition of two cycles of rapid bead beating (45 seconds at 4.5 m/second) using a FastPrep 24 homogeniser (MP Bio). Genomic DNA libraries were prepared using the TruSeq DNA v2 or TruSeq Nano DNA kit (Illumina), and whole genome sequencing was performed on an Illumina HiSeq 2500 at Medical Research Council Clinical Genomics Centre (Imperial College London) as previously described (36). All raw reads and information on lineages of isolates in this study have been submitted to the European Nucleotide Archive under the project accession PRJEB11842.

### Whole genome sequence analysis

Raw Illumina reads were aligned to the *Cn* reference genome H99 (37) using the Burrows-Wheeler Aligner (BWA) v0.75a mem algorithm (38) with default parameters to obtain high depth alignments (average 104x). Samtools (39) version 1.2 was used to sort and index resulting BAM files, and generate statistics regarding the quality of alignment. Picard version 1.72 was used to identify duplicate reads and assign correct read groups to BAM files. Furthermore, BAM files were locally realigned around insertions and deletions (INDELs) using GATK (40) version 3.4-‘RealignerTargetCreator’ and ‘IndelRealigner’, following best practice guidelines (41).

Single nucleotide polymorphisms (SNPs) and INDELs were called from all alignments using GATK (40) version 3.4-46 ‘HaplotypeCaller’ in haploid mode with a requirement that all variants called and emitted are above a phred-scale confidence threshold of 30. Both SNPs and INDELs were hard filtered due to a lack of training sets available for *Cn* by running VariantFiltration with parameters “DP < 5 || MQ < 40.0 || QD < 2.0 || FS > 60.0”; this expression ensured low confidence variants were filtered out if they met just one of the filter expression criteria. Resulting high-confidence variants were mapped to genes using VCF-annotator (Broad Institute, Cambridge, MA, USA) and the latest release (CNA3) of the Cn reference genome H99 and gene ontology.

Some isolates were suspected of having non-haploid genomes due to the high number of low confidence variants. For these isolates, ‘HaplotypeCaller’ was repeated in diploid mode.

The average (mean) coverage for each isolate were determined using GATK (40) version 3.4-46 ‘DepthOfCoverage’ under default settings. The *Cn* H99 (37) was again used as reference. In order to determine aneuploidy, whole-genome coverage data was normalised and regions displaying normalised coverage equal to 2 were deemed diploid events (likewise, normalised coverage equal to 3 were deemed triploid events, and so on), whereas normalised coverage equal to zero was deemed a deletion event.

### Susceptibility testing

The susceptibility testing of all relapse isolates were performed with the MICRONAUT-AM susceptibility testing system for yeast (Merlin) as recommended by the manufacturer. MICRONAUT-AM allows the determination of MICs of amphotericin B, flucytosine, fluconazole, voriconazole, posaconazole, itraconazole, micafungin, anidulafungin and caspofungin, and commercialises the well-established, but laborious, CLSI broth microdilution technique. Briefly, for each isolate five colonies were used to prepare a 0.5-McFarland-standard suspension in 0.9% NaCl. 1:20 dilution was prepared in 0.9% NaCl and 1:5 dilution was prepared in 11ml RPMI broth provided with the kit. 100μl AST indicator and 50μl Methylene blue solutions were mixed with the broth for manual susceptibility testing. The broth was then inoculated onto Merlin MICRONAUT 96-well testing plates (100ul/well) and incubated at 30°C for 72 hours. The lowest concentration of an antifungal agent with no detectable growth (MIC) was determined for each isolate based on fungal growth (pink) or no growth (blue). Obtained MICs were interpreted according to C. albicans EUCAST (Vers. 7.0 / 12-08-2014) values (42, 43).

### Phylogenetic analysis

Whole-genome SNPs were converted into relaxed interleaved Phylip format. Rapid bootstrap phylogenetic analysis using 500 bootstrap replicates was carried out on isolates in total (Table 1) using RAxML-HPC version 7.3.0 (44) as described in Abdolrasouli *et al*. (45): 35 isolates from this study in addition to 27 isolates (‘non-study’) were included to show the phylogenetic context of true relapse infections. These non-study isolates, whilst from a clinical source, were not recurrent isolates and were not isolated as part of the clinical trials described in the earlier Methods section. Resulting phylogenies were visualised in FigTree version 1.4. (http://tree.bio.ed.ac.uk/software/figtree/). The same process was completed for each chromosome individually for all 62 isolates, using 250 replicates in the rapid bootstrap analysis.

### Gene Ontology and KEGG pathway analysis

Non-synonymous (nsSNPs) mutations unique to each timepoint for each pair were assessed for significantly overrepresented gene ontology (GO) annotations and metabolic pathways. Briefly, genes found to contain a nsSNP mutation were interrogated for overrepresented Biological Process Ontology in the *Cn* H99 database. GO terms that were found to be associated with genes mapping to the InterPro domain database were transferred to GO associations, using a *p*-value cut-off of *p* < 0.05. For metabolic pathway enrichment in genes containing nsSNPs, genes were interrogated against the KEGG (46) pathway source for *Cn* H99, using a *p*-value cut-off of *p* < 0.05.

### Identifying sites under selection

BayeScan 2.01 (47) uses an outlier approach to identify candidate loci under natural selection. The method uses the allele frequencies that are characteristic of each population and estimates the posterior probabilities of a given locus under a model that includes selection and a neutral model. The programme then determines whether the model that includes selection better fits the data. This approach allows the simultaneous assessment of the influence of both balancing and purifying selection. Loci under balancing selection will present low F_ST_ values whereas high F_ST_ values reflect patterns of local adaptation (purifying selection) (48). Analysis was not undertaken for the VNII and VNB lineages due to low numbers of isolates, which would be insufficient to overcome the strong population structure. VNI Isolates at day 0 were assigned to a population and their associated relapse isolates constituted the second population. Analyses were conducted using the standard parameters including a 50,000 burn in period and 100,000 iterations. Several analyses were conducted varying the prior odds (from 10, 100 to 1,000) for the neutral model.

## Results

### Clinical and demographic information

The study included paired isolates from 17 patients, with a median age of 32 years (IQR 26-36) and median CD4 count at CM diagnosis of 22 (IQR 9-71) cells/μL. Six patients were male, 9 were female, with the gender of two patients unrecorded. The median time between initial and recurrence isolates was 115 days (minimum 55 days, maximum 409 days). In those for whom ART status was known, 2 of 16 (13%) patients were already on ART at the initial CM episode; 6 out of 15 (40%) patients had not started ART prior to CM recurrence.

Detailed clinical notes were available for the recurrent CM episode in 7 patients: two (CCTP52 and RCT9) had not attended follow up and never started ART prior to admission with recurrence – both died of the recurrent CM. One patient (CCTP32) had not been taking fluconazole for 2 weeks prior to recurrence. In four patients (CCTP27, CCTP50, RCT24, IFNR63) who were adherent to both ART and fluconazole at recurrence were assessed as having CM immune reconstitution inflammatory syndrome (CM-IRIS).

### Sequencing of paired samples isolated from patients infected with *Cn*

Prior to sequencing, multiple colonies from a subset of isolates included in this study were analysed using MLST to investigate whether a mixed infection was present in the original CSF extract. The results show that mixed infections were not present in 12 out of 17 Pairs included in this study. One Pair (Pair 7) was only tested once, and allele types (AT) were not sufficient to conclude whether sequence type (ST) 100 or 196 was present in both original and recurrent isolate. On two separate attempts, STs for Pairs and 17 could not be determined, reflecting a need for whole-genome sequencing (WGS) to characterise these Pairs. STs for Pairs 1 and 6 were inconclusive, and suggestive of a mixed infection present.

We recovered an average of 23.9 million reads from each isolate, with an average of 98.8% of reads mapped to the Cn H99 reference genome (37), and an average coverage of 104 +/-31.2 (standard deviation). To enable comparative studies and detect micro-evolutionary changes, precise variant-calling was needed; variants were identified and false positive low-confidence variants were filtered out to provide a set of high-confidence SNPs (see Materials and Methods). Full alignment, coverage and variant calling statistics are provided in Supplementary Materials Table 5.

Due to a high number of low-confidence SNPs filtered out in some isolates, which is suggestive of heterozygous SNPs, variant calling was re-run in diploid mode (see Methods) for all isolates in Pairs 3, 4, 5 and 17 (results in Supplementary Materials Table 6).

A high level of diversity was observed within the VNB lineage, resulting in long branch lengths (Figure 1). Although all VNB isolates were mapped to the *Cn* H99 reference (37), which is a VNI lineage isolate, we do not believe SNP numbers observed in the VNB lineage are a consequence of phylogenetic distance to the reference genome. This is because SNP determination revealed only 21.6% SNPs were common to the three VNB Pairs included in this study (Pairs 3, 12 and 17), highlighting the large amounts of genetic diversity seen within this lineage.

**Figure 1.**
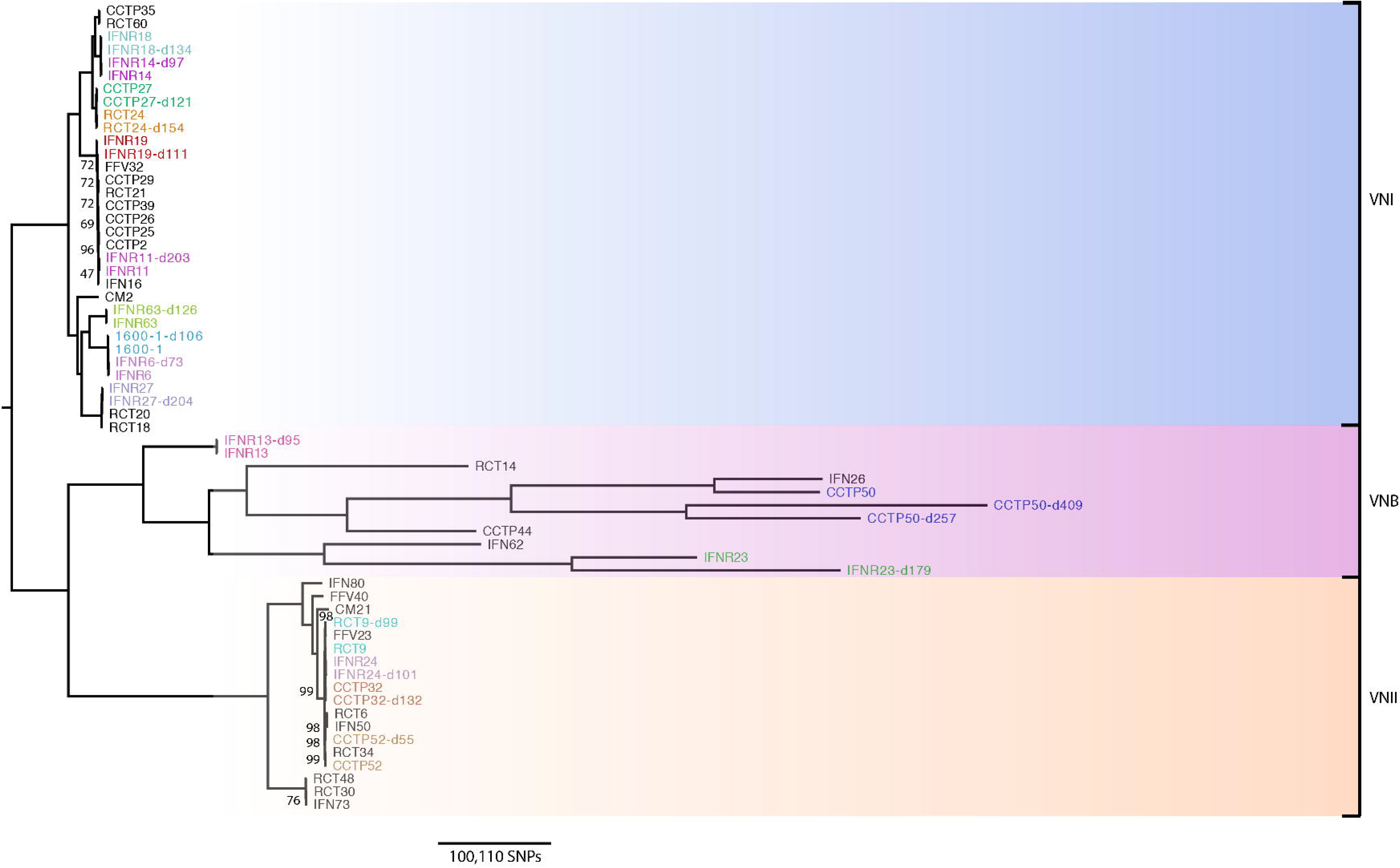
Phylogenetic analysis of *Cn* var. *grubii* isolates in this study (coloured), with additional isolates (shown in black) added to distinguish true relapse infections, or recurrent infections, and associated lineages. We hypothesise that isolates resulting from true relapse infections would be closely related phylogenetically. Bootstrap analysis over 500 replicates was performed on WGS SNP data from 62 isolates, including the 35 isolates included in this study, to generate an unrooted maximum-likelihood phylogeny, with all branches supported to 69% or higher (with the exception to a particularly clonal VNI clade, including Pair 15 only, which only had 47% branch support). Branch lengths represent the number of SNPs between taxa.

Phylogenetic analysis showed that of the 17 pairs of relapse isolates, three pairs were lineage VNB, whilst four and ten belonged to lineages VNII and VNI respectively (Figure 1). The average pairwise SNP diversity was far higher amongst isolates from the VNB lineage (140,835 SNPs) compared to isolates in the VNI (17,808 SNPs) and VNII (SNPs) lineages, showing that the VNI and VNII lineages are less diverse than VNB across our cohort. On average, isolates of the VNB, VNII and VNI lineages accumulated 365, and 3 unique SNPs per day between the time of the original isolation and the recurrence of infection. Isolates in the VNB lineage were more likely to experience a ploidy event, with an average of 1.6 changes in ploidy per isolate. Less than one isolate in the VNII and VNI lineages would, on average, experience ploidy events (0.375 and 0. respectively).

All pairs, with the exception of Pair 7, were isolated from patients in South Africa; Pair was isolated from a patient in Uganda. We classified the second isolate as a relapse of the original infection if more than 97% of SNPs were in common between original and recurrent isolates. The majority of pairs had >99% SNP similarity (Table 3) between original and recurrent isolates, with Pairs 6, 7 and 14 displaying 97%, 98% and 97% similarity respectively. Therefore, all pairs, with the exception of Pairs 3 and 17 (SNP similarity 44% and 56%) could be classified as relapsed infections on this basis. This confirms previous results obtained by MLST, and that the original and recurrent isolates sequenced of Pairs 1 and 7 (which had previously demonstrated a potential mixed infection) were indeed true relapse infections.

Within the VNB pairs (3, 12 and 17), the accumulation of SNPs between original and recurrent infection varied widely. We observed 178 and 304 SNPs/day for CCTP50-d and CCTP50-d409 respectively (Pair 3), 8 SNPs/day for Pair 12, and 968 SNPs/day for Pair 17. Due to the variation in SNP accumulation between Pair 12 and Pairs 3 and 17, we hypothesised that Pair 12 was a true relapse of the original infection, whilst Pairs 3 and 17 are showing inflated SNP numbers due to reinfection or an anomalous rate of evolution.

### Antifungal susceptibility testing

Fluconazole susceptibility testing (Table 1) using the Etest^®^ (bioMerieux) was carried out for 12 isolates (including three paired isolates) in this study by the accredited central Microbiology laboratory in Cape Town at the time of the clinical episode; five of these (CCTP27-d121 in Pair 1; CCTP50 and CCTP50-d257 in Pair 3; RCT24-d154 in Pair6; IFNR11-d203 in Pair 15) had MICs above the established epidemiological cut-off value (≥ 8μg/ml) for fluconazole (FLC). All pairs were retested following 5-10 years frozen storage in glycerol using the MICRONAUT-AM system for yeast susceptibility (Methods): all were found to be sensitive to FLC.

The fourfold increase in FLC MIC observed in Pairs 1 and 3 initial and recurrent infections provide a sound basis for relapse of infection due to drug resistance: in Pair (patient CCTP27), the initial isolate had a susceptible FLC MIC of 4 ug/ml, whilst the recurrent isolate was resistant at an MIC of 64 ug/ml; in Pair 3 (CCTP50), the initial isolate MIC was 16 ug/ml (intermediate), whilst a highly resistant MIC of 256 ug/ml was found on recurrence at day 257.

### Serial isolates share a recent common ancestor, suggesting relapse of infection

To investigate whether the Cn isolated from the same patient were relapse infection of the original isolate, or infection with a new isolate, we undertook phylogenetic analyses to determine their relationships.

As described above, the high level of common SNPs, and subsequent low level of unique SNPs, between recurrent isolates indicated that all pairs, with the exception of Pairs and 17, were relapse of the original infections (Table 3). Phylogenetic analysis (Figure 1) confirmed that all pairs (excepting Pairs 3 and 17) clustered together with short branch lengths, confirming the low level of divergence between original and recurrent isolates, thus confirming that they were relapse of the original infections. However, only 46% and 56% of SNPs were found to be in common between initial and relapse infection in Pairs 3 and 17 respectively (Figure 1 and Table 3). These VNB pairs (Pair 3; CCTP50, CCTP50-d257 and CCTP50-d409 and Pair 17; IFNR23 and IFNR23-d179) showed markedly longer branch lengths, suggesting either reinfection or elevated rates of within-host evolution. Further analysis was undertaken to confirm or refute that reinfection by a different isolate was responsible for Pairs 3 and 17. Phylogenetic analysis for all isolates included in Figure 1 were repeated for each of the 14 *Cn* chromosomes individually (Supplementary Figure 1).

Phylogenetic analysis of Pair 3 showed that the original infecting genotype of CCTP was highly related to the isolate IFN26 (not included in this study, but included in the phylogeny to assist with defining lineages - see Materials and Methods). All three genotypes from Pair 3 were found to be phylogenetically clustered together, but with long branches (Figure 1). Chromosome-by-chromosome analysis indicated that Pair serially isolated genotypes displayed differing relationships for each chromosome, and all three serial genotypes were clustered together only in the phylogeny for chromosome 1 (Supplementary Figure 1). All three genotypes were phylogenetically similar for three other chromosomes, however long branches and clustering with additional non-study isolates suggested differing evolutionary relationships. The three serially isolated genotypes of Pair 3 were completely phylogenetically dissimilar in three chromosomes; the remaining chromosomes saw either the day 1 isolate (CCTP50) and day 409 isolate (CCTP50-d409), or the day 257 isolate (CCTP50-d257) and day isolate (CCTP50-d409) phylogenetically more related.

Pair 17 isolates (ID IFNR23) clustered together in only two of the 14 chromosomal phylogenies explored (Chromosomes 10 and 12); in the remaining chromosomal phylogenies, the Pair 17 isolates either displayed a close phylogenetic relationship, but with long branches (6 chromosomes), or were phylogenetically distinct from one another, and were more phylogenetically related with other study or additional isolates (6 chromosomes).

### Microevolution within the human host

Our data present a unique opportunity to observe microevolution of all three lineages of *Cn* var. *grubii* in the human host. Although multiple factors determine evolutionary rates, identifying non-synonymous SNPs (nsSNPs) that cause amino acid change is a standard method for inferring genetic diversity and observing natural selection on codons.

Less than 3% of nsSNPs were unique to recurrent isolates in all pairs, further suggesting that all pairs are relapse of the original infection, with the exception of the VNB Pairs and 17 (15.2% and 59.8% of all nsSNPs are unique, respectively).

SNPs unique to each timepoint for each pair were identified. All SNPs at Day 1 in all pairs, and all SNPs at time of isolate of recurrent infection in all pairs, were compared. No SNPs were found to be common to all 17 pairs at either Day 1 or at point of recurrent infection; however, there were VNII and VNB lineage-specific, and timepoint-specific, common SNPs.

Five SNPs, all intergenic, were found to be common at Day 1, along with five different SNPs, also intergenic, within VNB pairs (Pairs 3, 12 and 17). Three intergenic SNPs were common to all VNII pairs (Pairs 2, 4, 5 and 9) at timepoint Day 1, whilst 14 SNPs were common to all VNII pairs at the point of recurrent infection, five of which were intergenic. The remaining 9 SNPs were located in the 5' untranslated region (UTR) gene *SMF1* (CNAG_05640), a metal ion transporter with a natural resistance-associated macrophage protein. Selection analysis indicated that this gene was not under selection pressure, however.

To evaluate the genetic divergence, Wright's fixation indexes (*F_ST_*) were calculated to identify SNPs under selection in VNI original and recurrent infection populations investigating 96,856 loci (see Methods). No putative loci under either diversifying or balancing selection could be detected using a false discovery rate (FDR) of 0.05. *F_ST_* values were limited to not exceed 3.47^×010-5^.

### Aneuploidy as a generator of diversity in recurrent infection

Normalised whole-genome coverage was plotted to observe possible aneuploidy (increase or decrease in copies of chromosomes) and copy number variation (CNV) events. Aneuploidy events were observed in 7 genome pairs, suggesting either interspersed or tandem duplications of large segments of the genome.

Ormerod *et al*. (5) previously published a study showing relapse isolates exhibiting aneuploidies of chromosome 12. We observed aneuploidy of chromosome 12 in four pairs (Pairs 1, 5, 10 and 14) included in this study. The aneuploidy spanned different regions of chromosome 12 in all pairs, but all aneuploidies were present in the right arm of the chromosome: in Pair 5 the aneuploidy was restricted to 392 kbp of one chromosome arm; in Pair 1, the aneuploidy spanned an entire arm of chromosome (603 kbp). Pair 14 displayed this aneuploidy in both the initial and recurrent infection, and the aneuploidy spanned the whole chromosome; the recurrent infection isolate of Pair 10 also displayed aneuploidy along the whole chromosome. Evaluation of the read depth along chromosome 12 revealed triplication of the chromosome 12 arm in the Pair 1 recurrent infection isolate, a phenomenon also seen in Ormerod *et al*. (5). Further analysis of read depth in the Pair 5 recurrent isolate revealed a diploid genome, and chromosome 12 was also experiencing triploidy. Current annotation of the *Cn* H genome reveals the presence of 327 genes in chromosome 12; the right arm of chromosome 12 has 260 genes present. We scanned the genes present in this arm of chromosome 12 for genes potentially involved in virulence, which might prove advantageous to the progression of infection or drug resistance. One such gene was *SFB2* (CNAG_06093), which is involved in the conservation of the sterol regulatory element-binding protein pathway (SREBP) (49). An alcohol dehydrogenase (*GNO1* – CNAG_06168) was also present, which is thought to be involved in the defence against host response (50). Analysis for enrichment of metabolic pathways also revealed that the genes present in this chromosome arm are significantly involved in the metabolism -2 of drugs (corrected *p*-value *p* < 3.81e^-2^).

We searched for aneuploidies in genes known to be involved in drug resistance and virulence. *CAP10* appeared to be haploid in all isolates, with the exception of Pairs and 5, where the initial infection (CCTP52) and recurrent infection (RCT9-d99) were found to be diploid, respectively. However, on closer inspection (see Methods), we believe that the isolates in Pairs 4 and 5 (CCTP52 and RCT9-d99) have diploid genomes, implying that the *CAP10* gene is actually tetraploid. Whether the remaining isolates in these two pairs (CCTP52-d55 and RCT9) have diploid genomes could not be distinguished, however, it is clear that *CAP10* loses ploidy from initial infection to relapse for Pair 4, with no evidence of loss of heterozygosity (LoH), whilst the reverse is true for Pair 5. *CAP10* was also found to be tetraploid (as the genomes of these isolates were found to be diploid) in both initial and recurrent infections for Pairs 3 and 17.

The *ERG11* gene on chromosome 1 was found to have increased copy number in numerous pairs (2, 3, 4, 5, 9, 12, and 17), and was not found to be lineage-associated. However this CNV was maintained throughout infection to recurrence in all pairs, with the exception to Pair 4; since Pair 4 initial infection (CCTP52) was found to have a diploid genome, *ERG11* was tetraploid, and lost this ploidy to be diploid with respect to the rest of the genome in the recurrent infection (CCTP52-d55). Whilst chromosome 1 was duplicated in the initial infection isolate of Pair 15 (IFNR11), *ERG11* was found to be haploid; the ploidy of chromosome was subsequently lost in the recurrent infection isolate of Pair 15 (IFNR11-d203).

*ERG11* in Pair 4 (ID CCTP52) did not have any nsSNPs in the original infection (CCTP52), but one nsSNPs was present in *ERG11* in the recurrent infection (CCTP52-d55). Clinical notes show that the patient from which Pair 4 was isolated was given fluconazole (mg/d) on initial infection, did not attend follow up or receive ART or further fluconazole, and was then re-admitted and died from CM recurrence at day 55. MIC values were unfortunately not available for either original or recurrent isolates.

### Nonsense mutations in DNA mismatch repair genes cause hypermutator states

Phylogenetic analysis on a chromosome-by-chromosome basis revealed that Pair isolates only clustered together in two of the fourteen chromosomes; the three isolates were phylogenetically dissimilar in 4 chromosomes, whilst day 257 and 409 isolates (Pair 3 CCTP50-d257 and CCTP50-d409) were phylogenetically more similar to each other than to the day 1 isolate (CCTP50) in five chromosomes. Day 1 and day 409 isolates were more phylogenetically similar than to the day 257 isolate in three chromosomes. The lack of phylogenetic similarity (12 out of 14 chromosomes) shown in the three isolates of Pair 3 indicated that these three isolates do not show a recent common ancestor, and provides evidence for reinfection with a new isolate, rather than relapse. In contrast, the two isolates in Pair 17 were only phylogenetically related for five out of the 14 chromosomes (Supplementary Figure 1), suggesting that on this basis the recurrent infection was distinct enough as to be defined as a non-relapse infection in this Pair. However, on further investigation this was found not to be the case.

We analysed the coverage profiles and synonymous/non-synonymous ratios of both isolates in Pair 17 (Figure 2c). Although the aneuploidies observed were extensive throughout the genome, the increases in ploidy appeared on similar chromosomes in both isolates. A similar observation was seen for the strikingly increased number of nsSNPs in both isolates in Pair 17: 41,549 and 46,622 nsSNPs for IFNR23 and IFNR23-d179 respectively; an even larger increase in synonymous SNPs was also observed (68,094 and 82,172 synonymous SNPs for IFNR23 and IFNR23-d179 respectively). We therefore sought to identify a mechanism responsible for the high number of synonymous and nsSNPs, and ploidy.

**Figure 2.**
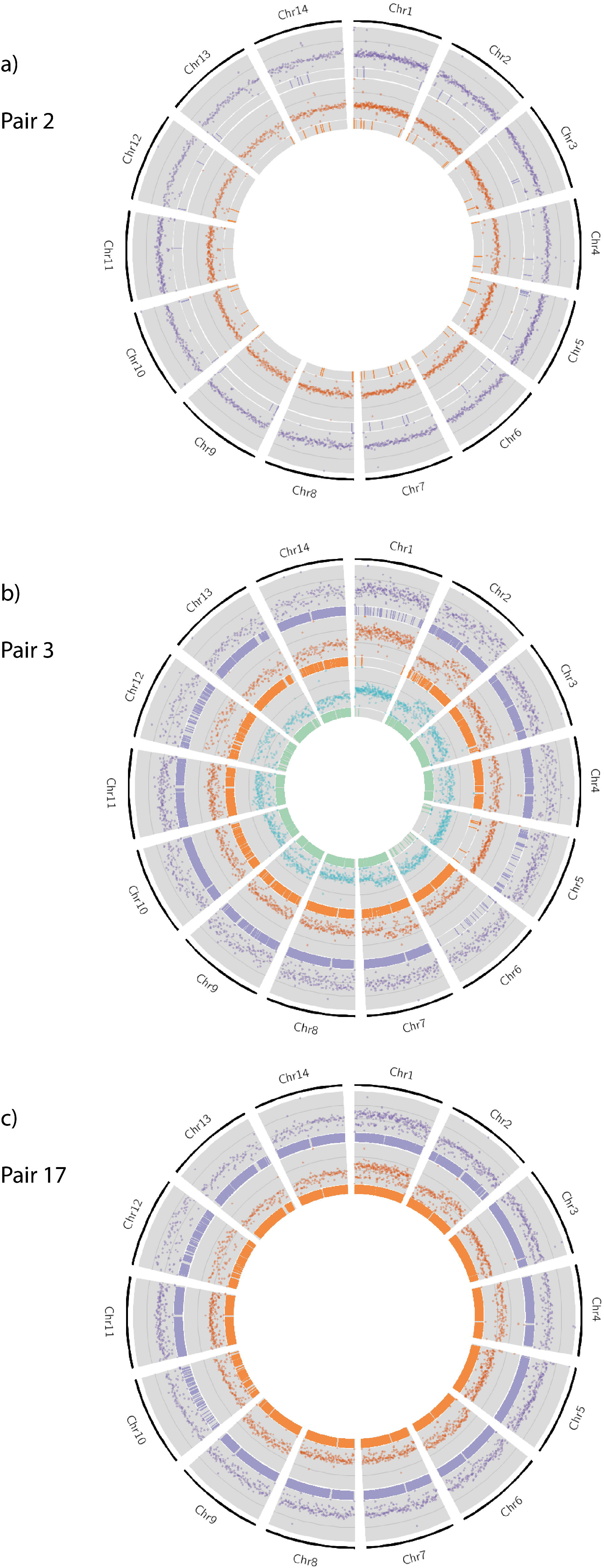
Extensive chromosomal copy number variation was observed in all isolates in Pairs 3 and 17, when compared to H99. Pair 2 is included to illustrate isolates without ploidy and extensive nsSNPs. Here, normalised whole-genome depth of coverage is shown, averaged over 10,000-bp bins, in scatter plots. Bar plots represent the position of nsSNPs. The purple track represents the original isolate, orange the recurrent isolate, and green (in the case of Pair 3) for the final recurrent isolate. a) No increase in ploidy is observed in either the original or recurrent isolate of Pair 2, and a small number of nsSNPs are seen. b) Increase in ploidy is observed in many chromosomes in the Day isolate for Pair 3, some of which are lost over time. A large number of nsSNPs are observed in all chromosomes in isolates of Pair 3, with chromosome 6 being the exception: very few nsSNPs are located in chromosome 6 in CCTP50 and CCTP50-d409, whereas over 2000 nsSNPs are observed in chromosome 6 in CCTP50-d257. c) a gain in ploidy is observed for Chromsomes 2, 4, 6 and 9 compared to the Day 1 isolate in Pair 17, whereas ploidy remains unchanged for Chromosomes 1 and 12.

Previous studies have reported that mutations in the DNA mismatch repair gene *MSH2* have resulted hypermutator effects in bacteria and the yeast *S. cerevisiae* (51). Both Pair 17 isolates were found to harbour two nonsense (i.e. point mutations in the DNA sequence that result in a premature stop codon) mutations within the coding region of the gene encoding *MSH2*, the DNA mismatch repair protein. Nonsense mutations in *MSH2* were not observed in any other pairs included in this study. These mutations were in the same positions in both the original and recurrent isolates (Ser-888-STOP and Ser-86-STOP).

We then performed a genome-wide search in both Pair 17 isolates to identify further nonsense mutations in DNA mismatch repair genes. Both Pair 17 isolates were found to harbour a single nonsense mutation within the coding regions of genes encoding *MSH5* and *RAD5*. Again, nonsense mutations in these genes were not observed in any other pairs included in this study. These nonsense mutations caused Gln-1066-STOP in *RAD5*, and Gln-709-STOP in *MSH5* in both original and recurrent isolates.

Since the likelihood of such mutations occurring by chance in independent genomes lacking a common ancestor is very small, this suggest that rather than being a reinfection, this was indeed a relapse of the original infection, and the phylogenetic dissimilarity between the two isolates was due to hypermutation. A total of 293 SNPs were located in *MSH2* in both Pair 17 isolates, compared to a mean of 30 SNPs per isolate in the remaining Pairs that we studied (Supplementary Table 2). More SNPs overall were observed in the recurrent isolate (IFNR23-d179) in both *RAD5* and *MSH5* (361 and 357 respectively) when compared to the original isolate (IFNR23 – 320 for *RAD5*, 305 for *MSH5*). These numbers are considerably higher than the average of SNPs and 37 SNPs per isolate in the remaining Pairs included in this study for *RAD5* and *MSH5* respectively.

## Discussion

Relapse of CM caused by *Cn* is usually due to the persistence and recurrence of the original infecting isolate (52), and studies often focus on rates of within-host microevolution between serially collected isolates. However, recent studies have shown that an infection of a population of dissimilar genotypes is responsible for 20% of relapse cases (53). We used whole-genome sequencing to distinguish co-infections of a population of genotypes from a relapse of a single genotype owing to treatment failure (Figure 3). Using WGS we can distinguish between relapses of infection of the same genotype, which differ by only a few SNPs, whilst initial infection by a population of dissimilar genotypes will see a difference of many SNPs between initial and relapse infection as genetic drift occurs. Our results show that *Cn* incurs numerous unique small and large-scale changes during infection, and that a subset of these may have adaptive value. Whilst this study is concerned with the genomics of recurrent infections by identifying SNP changes and ploidy potentially involved in the persistence of *Cn* infection, future work should investigate the potential role of gene expression changes and gene networks involved in changes in fitness amongst the populations of infecting genotypes that underpin the recurrence of infection. Previous studies in *Brucella* infection and TB have highlighted the merits of using transcriptomics to identify patients requiring more intensive treatment (54) and differentiating between dormancy and reactivation (55) respectively. Such approaches are likely to increase our understanding of clinical cases of recurrent *Cn* infections through identifying the genetic basis of phenotypic switching (56) and the gene regulatory networks involved in latency, virulence and resistance to antifungal therapies.

**Figure 3.**
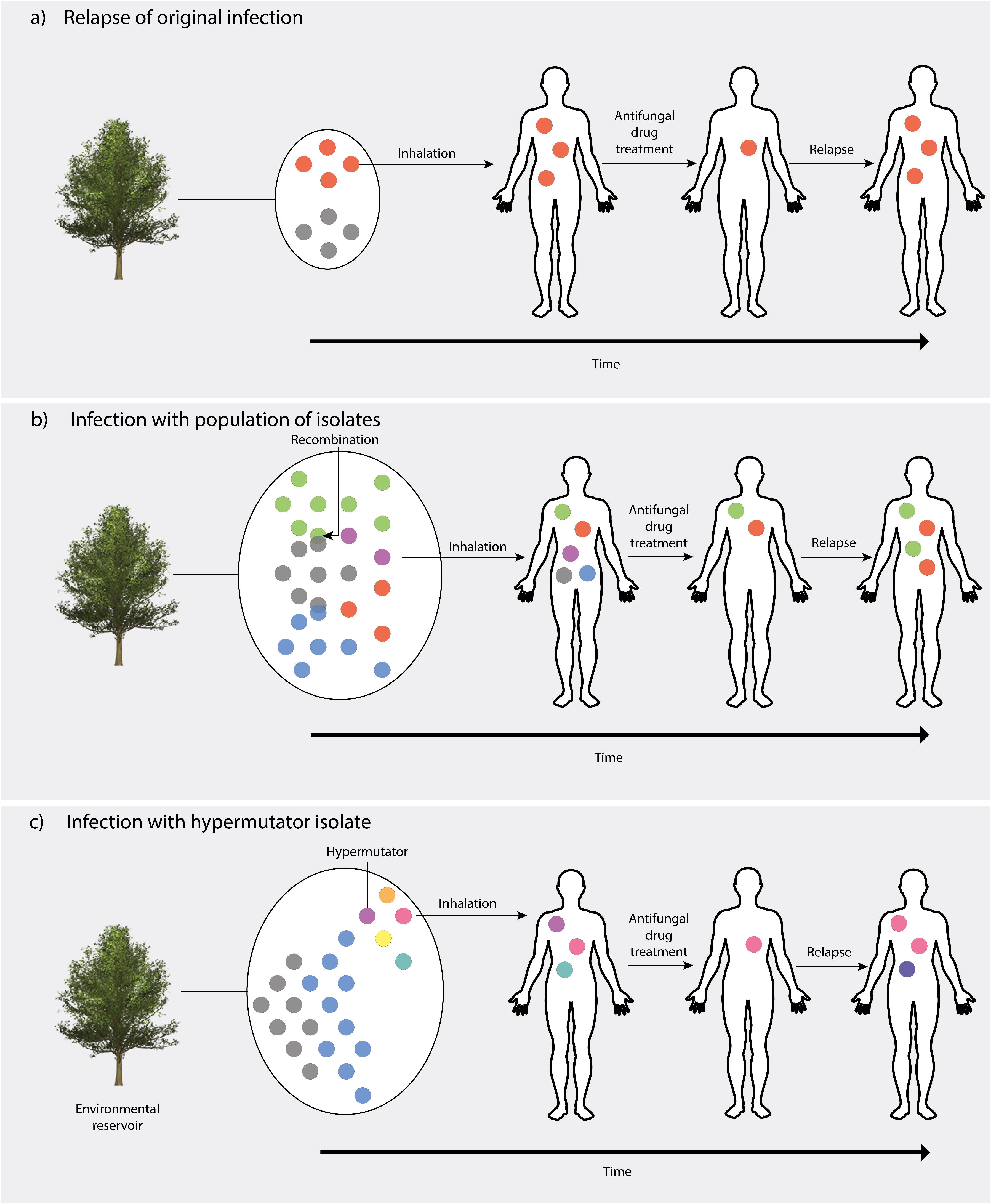
Hypotheses of routes of infection of the human host by *Cn* var. *grubii*. a) Inhalation of a single population of basidiospores into a new host. Due to low within-host diversity and being drug naïve, there will be a bottleneck in population size due to antifungal drug treatment. However, if the initial drug regimen is insufficient to sterile the CSF, resistance may develop on FLC maintenance therapy due to selection pressure, resulting in relapsed infection from proliferation of a drug-resistant isolate. b) VNB lineage *Cn* exists the environment as a population, which can undergo recombination to produce genetically similar isolates, but with significantly diversity. Due to transmission bottlenecks, only a sample of the pathogen diversity will be transferred to the host, in this case, by inhalation, but it is possible for a population of *Cn* to infect a single immunocompromised individual. Some isolates may be susceptible to antifungal drugs and are thus becoming removed, whilst other isolates may be inherently resistant and hence cause a relapse infection. c) Mutations in the DNA mismatch repair gene *MSH2* cause an isolate to become a hypermutator. Some genotypes may be susceptible to antifungal drugs, but the high mutation rate allows the infection to adapt rapidly to the host and evolve drug resistance. These genotype proliferate in the host, thus causing relapse infection.

One pair (Pair 3) included in this study did not display a relapse of the initial infecting isolate. Analysis of this pair showed that only 46% of SNPs were in common between the initial and recurrent infection, suggesting that relapse was caused by a new, albeit similar, genotype. The extensive chromosomal copy number variations, or aneuploidies, observed in Pair 3 (Figure 2) also show that different genotypes were isolated at subsequent timepoints (days 257 and 409). Phylogenetic analyses showed that this pair belongs to the VNB lineage; it is known that a population of VNB genotypes can be found in one location, such as on the same tree (57). Therefore, it is possible for a single immunocompromised individual to inhale a cluster of basidiospores from a single mating population, which would lead to a cluster of related, but recombined genotypes that then come to dominate the infection at different timepoints. Although a population can reside in an environmental reservoir, recombination between genotypes can occur, generating closely related, yet distinct, genotypes (Figure 3b). This latter hypothesis supports our observations of differing numbers of nsSNPs between day (CCTP50-d257) and the original (CCTP50) and day 409 (CCTP50-d409) isolates, as well as the ploidies and MIC values (Table 1) seen at the time of sample isolation: the day isolate (CCTP50) initially had an intermediate FLC MIC of 16 ug/ml, whilst the recurrent isolate at day 257 had a highly resistant FLC MIC of 256 ug/ml. These MIC values are suggestive of drug-resistant genotypes being present and selected for within this patient by the prolonged maintenance on FLC monotherapy following induction therapy with amphotericin B. It is also likely that the population of VNB isolates circulating in the patient were not sufficiently sampled by sequencing only one colony at each timepoint, and that deeper sequencing would have uncovered greater genomic diversity.

The occurrence of aneuploidy is seen as an evolutionary process that rapidly alters fitness, and has been described in multiple human fungal pathogens as a means of generating drug resistance (6, 58). Sionov *et al.* (6) reported the duplication of multiple chromosomes in response to high concentrations of FLC, which resulted in genotypes developing FLC drug resistance. Associated gene duplications in *Cn* chromosome included *ERG11* and *AFR1*, which are both transporters of azole drugs. Whilst duplications of *ERG11* were seen in seven pairs (2, 3, 4, 5, 9, 12 and 17), these were not necessarily associated with an entire duplication of chromosome 1. Sionov *et al.* suggested that *ERG11* contributed to the duplication of chromosome 1 (6); we observed only one isolate (IFNR11 of Pair 15) displaying a duplication of chromosome 1, but a single copy of *ERG11*, suggesting ploidy was not complete throughout the chromosome. Since this isolate was the initial infection, we can assume that the duplication of chromosome 1 was not solely due to stress of azole drug treatment, suggesting that ploidy can be activated under different conditions, such as the stress associated with adaptation to the host. A possible limitation is that the observed duplication may be due to prolonged frozen storage.

Ormerod *et al.* (5) showed an aneuploidy (duplication) in chromosome 12 between serially collected isolates. Four pairs included in this study (Pair 1, 5, 10 and 14) all showed aneuploidy in chromosome 12; however, Pair 14 (ID IFNR19) displayed this aneuploidy in both the initial and relapse infections. Since aneuploidies are typically lost upon removal of drug pressure (6), one can assume that this aneuploidy was maintained due to previous drug exposure potentially not reported by the patient, or that aneuploidy helps *Cn* adapt to the host environment (59). Chromosome 12 experienced triploidy in the Pair 1 recurrent isolate (CCTP27-d121); this pair also demonstrated drug resistance to fluconazole, with a FLC MIC of 4 at initial infection, and a FLC MIC of 64 at recurrent infection. Ormerod *et al.* (5) hypothesise that the large number of genes affected by the increased copy number of chromosome 12 contributes to metabolome differences; however, we hypothesise copy number variation of chromosome 12 is a response to FLC stress, resulting in increased MIC, and that some genes present on chromosome 12, such as *ERG8* and *CAP6*, may be targets of azole drugs or involved in *Cn* virulence.

Antimicrobial drugs impose strong selection pressure on pathogens, with may lead to the evolution of drug resistance (60, 61); there are, however, fitness costs associated with the evolution of resistance to antifungal drugs that may impact fitness (17). Genome-wide scans for sites under selection leads to the identification of possible sites of drug resistance. We did not identify any significant sites when comparing VNI original infection versus recurrent infection, and the number of VNB and VNII isolates were too low for analysis. Whilst these results could be interpreted as there being no sites under selection in the VNI isolates sampled in this study, it is more likely that similar patterns would not be seen amongst individuals due to stochasticity and clonal interference (61). It is also likely that as there is little recombination in VNI isolates compared to VNB and VNII isolates (62-64) and therefore linkage is complete across the genome, further hampering selection analysis. We therefore found no evidence for genetically determined alterations in drug resistance in the study isolates.

MIC values were only obtained for 9 out of 35 isolates in this study at the time of sampling. Susceptibility testing at a later date revealed all the isolates to be susceptible to antifungal drugs including FLC, suggesting that any resistant phenotypes had been lost in the absence of drug selective pressure. It is therefore important for clinicians to request susceptibility testing in real time, at the very least in all cases or recurrent CM.

Whilst Pair 17 did not exhibit a high percentage of common SNPs between the original and recurrent isolates indicative of a relapse infection, the elevated rate of SNPs observed in all chromosomes of both isolates suggested this was not a re-infection as seen in Pair 3 (Figure 3c). Rather, our results suggest that the isolates in this pair were exhibiting a hypermutator phenotype, as a result of two nonsense mutations in the DNA mismatch repair gene *MSH2*, and one nonsense mutation in each of the DNA mismatch repair genes *RAD5* and *MSH5*. Whilst previous studies have shown hypermutator phenotypes aid adaptation to stress (65), and we hypothesise that hypermutation may lead to adaptation of drug resistance under the stress of antifungal treatment. These results are the first to the authors' knowledge to report on nonsense mutations in *MSH2, RAD5* and *MSH5* in *Cn*. Further investigation is required to determine whether these nonsense mutations have a role in drug resistance phenotypes using transcriptomic approaches and creating single-gene knockout mutants of *MSH2, RAD5* and *MSH5*. It is also necessary to test the virulence of hypermutator isolates in the mouse model and to describe the impact of the increase in mutation rate that occur a result of this hypermutation. Our study only includes one pair of hypermutator genotypes, so further sampling is required to identify whether this phenomenon is specific to the VNB lineage, whether hypermutators occur in the VNII and VNI lineages, and whether they are clinically relevant.

This work represents the most extensive comparative genome-sequencing based study to investigate microevolution in serially collected isolates of *Cn* to date. The observation of an infection of a single patient with a population of VNB isolates is clinically relevant, as widely used drug regimens with azole monotherapy may not be effective against such a genetically diverse infection. It is also likely that the extensive genetic diversity seen in clinically isolated VNB isolates may be due to mixed infection. Hypermutation due to nonsense mutations in the DNA mismatch repair genes *MSH2, RAD5* and *MSH5* cause an increased mutation and rate of aneuploidy in *Cn*, which may confer an increased ability to adapt to drug pressure. Further sampling is required to identify whether hypermutation is a phenomenon only observed in the VNB lineage, and how these mutations impact the fitness of *Cn* by imposing a high genetic load.

## Acknowledgements

Special thanks to Mr. Ian Wright for his financial contribution to this project, and Dr. Winnie Wu and StarLab UK for supplying filter pipette tips free of charge. This work was funded by a Medical Research Council grant (MRC MR/K000373/1) awarded to MCF and a grant from Imperial College London (WPIA/F24085) awarded to JR. JJ was supported by the Wellcome Trust (training fellowship, WT081794) and is supported by a grant from the Penn Center for AIDS Research (CFAR), an NIH-funded program (P30 AI 045008). Thankyou to Sean Lobo and Ahsan Awais Bokhair for initial work on MICs and MLST, respectively.

## Author contributions

TB, JJ, JS, AR, GM and TSH conducted the clinical studies and provided isolates included in this study. TB, JJ and MCF conceived the research question. JR and MB conceived the experiments; MB performed library preparations for sequencing, with assistance from MV. JR performed alignments, variant calling and downstream analysis, and wrote the paper. MV performed BayeScan analysis. All authors read and contributed to this paper.

## Supplementary material

Supplementary Table 1 - MLST results of independently testing three colonies per study isolate.

Supplementary Table 2 - Number of SNPs in *MSH2* in each isolate, categorised by type or location. SYN = synonymous SNP, NSY = non-synonymous SNP, p5UTR = 5' UTR, p3UTR = 3' UTR, NON = nonsense mutation.

Supplementary Table 3 - Non-synonymous SNPs in Pair 3 isolates, per chromosome

Supplementary Table 4 - Number of non-synonymous and synonymous SNPs per isolate

Supplementary Table 5 - Details of alignment and variant calling for all paired isolates included in this study

Supplementary Table 6 - Number of homozygous and heterozygous SNPs in isolates of Pairs suspected of diploidy.

Supplementary Figure 1 – Phylogenetic analysis each chromosome of *Cn* var. *grubii* isolates in this study (coloured) with additional isolates (shown in black), to identify whether Pair 3 consists of different isolates or is a relapsed infection. We hypothesised that if Pair 3 was a relapsed infection, all isolates would share the same phylogenetic relationship in all 14 chromosomes. Rapid bootstrap analysis over 250 replicates was performed on SNP data from 62 isolates to generate an unrooted maximum-likelihood phylogeny (only branches not supported to 100% are indicated above branches). Branch lengths represent the number of SNPs between taxa.

Supplementary Figure 2 – Chromosomal copy number variation can be observed in some isolates in Pairs 1, 4, 5, 10, 14 and 15, when compared to the *Cn* reference genome, H99. Normalised whole-genome depth of coverage is illustrated here, averaged over 10,000-bp bins and represented as a scatter plot. Bar plots represent the position of nsSNPs, where the purple track represents the original isolate, and orange represents the recurrent isolate. Note that Pairs 2, 3 and 17 are not present in this Figure, as they are represented in Figure 2.

